# Primary macrophages exhibit a modest inflammatory response early in SARS-CoV-2 infection

**DOI:** 10.1101/2022.02.02.478897

**Authors:** Ziyun Zhang, Rebecca Penn, Wendy S Barclay, Efstathios S Giotis

## Abstract

Involvement of macrophages in the SARS-CoV-2-associated cytokine storm, the excessive secretion of inflammatory/anti-viral factors leading to the acute respiratory distress syndrome (ARDS) in COVID-19 patients, is unclear. In this study, we sought to characterize the interplay between the virus and primary human monocyte-derived macrophages (MDM). MDM were stimulated with recombinant IFN-α and/or infected with either live or UV-inactivated SARS-CoV-2 or with two reassortant influenza viruses containing external genes from the H1N1 PR8 strain and heterologous internal genes from a highly pathogenic avian H5N1 or a low pathogenic human seasonal H1N1 strain. Virus replication was monitored by qRT-PCR for the *E* viral gene for SARS-CoV-2 or *M* gene for influenza and TCID_50_ or plaque assay, and cytokine levels were assessed semiquantitatively with qRT-PCR and a proteome cytokine array. We report that MDM are not susceptible to SARS-CoV-2 whereas both influenza viruses replicated in MDM, albeit abortively. We observed a modest cytokine response in SARS-CoV-2 infected MDM with notable absence of IFN-β induction, which was instead strongly induced by the influenza viruses. Pre-treatment of MDM with IFN-α enhanced proinflammatory cytokine expression upon infection. Together, the findings concur that the hyperinflammation observed in SARS-CoV-2 infection is not driven by macrophages.

## Introduction

The ongoing COVID-19 pandemic has generated many urgent questions on the diverse clinical manifestations of the causative agent Severe Acute Respiratory Syndrome Coronavirus 2 (SARS-CoV-2). SARS-CoV-2 requires the binding of its spike surface protein to a cellular receptor, the angiotensin-converting enzyme 2 (ACE2), to gain entry into host cells and activation by a host cell proteases TMPRSS2 [1–3]. The variable expression of ACE2 in different tissues across individuals and polymorphisms in both ACE2 and TMPRSS2 genes contribute to COVID-19 severity/fatality variations [4–7]. However, the severity of the disease outcome is widely believed to be associated with a derangement of the immune system such as a delayed type I/III interferon response and underlying co-morbidities in infected patients [8, 9]. One of the perplexing hallmarks of SARS-CoV-2 infection is the exacerbated inflammatory response in severe COVID-19 patients resulting in excessive release of pro-inflammatory cytokines known as “cytokine storm”, leading in turn to detrimental alveolar damage and fibrosis, progressive respiratory failure and multiple organ dysfunction [10]. A similar excessive inflammatory reaction has been observed in other zoonotic respiratory viruses such as SARS and MERS coronaviruses as well as human infections with avian influenza viruses such as H5N1 [11–13] whereas seasonal influenza induces a less severe response [14]. The prototypical influenza virus-induced cytokine storm has been described to originate from several cell types such as tissue macrophages, mast, endothelial, and epithelial cells [15]. These cells upon virus stimulation release initially TNF-α and IL-1ß, which in-turn stimulate the release of other cytokines mainly IL-1, IL-6, IL-8 and macrophage inflammatory protein-1α (MIP-1α) [13, 15, 16]. Although both influenza and COVID-19 are associated with hyperinflammation, there are marked differences between the two conditions in respect of origin, biochemical abnormalities and pathophysiology [17]. Nonetheless, numerous studies have positively correlated the elevated plasma levels of key proinflammatory cytokines in patients with disease severity and mortality for both COVID-19 and highly pathogenic influenza [13–15, 18, 19].

Understanding the precise drivers of SARS-CoV-2-induced hyperinflammation and their correlation to disease outcome is crucial to guide targeted therapeutic interventions. The degree to which SARS-CoV-2 targets the diverse cytokine-producing cells (*i.e*. macrophages, B and T lymphocytes, mast cells, endothelial cells, fibroblasts and various stromal cells) has not yet been fully elucidated; hence, their individual role in initiating, contributing and/or sustaining the cytokine storm remains unclear. A transcriptomic study reported that SARS-CoV-2 sequencing reads were detected in COVID-19 patients’ peripheral blood mononuclear cells [20], suggesting that SARS-CoV-2 may be able to replicate in specific immune cell subsets. Infection of these cell subsets may explain the persistence of the virus after pneumonia is resolved in some COVID-19 patients [21]. Macrophages, in particular, are critical for activation and resolution of systemic inflammation and are rapid producers of both proinflammatory and regulatory cytokines in response to local inflammation and pathogen infection [22]. Post mortem analyses and sequencing approaches have revealed that the lungs of COVID-19 patients with severe disease are infiltrated with macrophages suggesting they play a key role in COVID-19 pathophysiology [23–25]. Earlier reports suggested that alveolar macrophages are infected by SARS-CoV-2 [24–26] and that the diverse expression of ACE2 on macrophages among individuals might govern the severity of SARS-CoV-2 infection [27]. Since then, several studies have demonstrated that macrophages are not permissive to SARS-CoV-2 infection [28–31]. Conversely, some reports argue that macrophages orchestrate the cytokine storm [21, 24, 25, 30, 32] while other propose that macrophages play a secondary role in the virus-associated inflammation [29, 31].

Here, we report that primary macrophages are refractory to SARS-CoV-2 and induce modest levels of pro-inflammatory cytokines upon SARS-CoV-2 infection compared to two influenza strains. Pre-exposure of macrophages to exogenous IFN-α exacerbated the virus-induced inflammatory response. These results concur with the findings of previous reports that macrophages are not the original source of pro-inflammatory cytokines early during infection.

## Materials and methods

### Cells and viruses

Human macrophages were purchased from Lonza Group AG (Switzerland) and were generated in the presence of human macrophage colony-stimulating factor (hM-CSF) from CD14^+^ human monocytes. Monocytes derived from a 54-year-old, HBV-, HCV- and HIV-negative, African American male. The macrophages were seeded 5×10^5^ (0.5 ml) per well, in 24-well plates, with X-Vivo 15 media (Lonza Group AG), 25 ng/μl of hM-CSF (Gibco, USA) and 10% Fetal Bovine Serum (Gibco, UK). African green monkey kidney cells (Vero E6; ATCC CRL-1586) were maintained in DMEM, 10% FCS, 1% non-essential amino acids (NEAA) and 1% penicillin-streptomycin (P/S). Human epithelial colorectal adenocarcinoma cells (Caco-2; ATCC HTB-37) and human lung cancer cells (Calu-3; ATCC HTB-55) were maintained in DMEM, 20% FCS, 1% NEAA and 1% P/S. All cell cultures in this study were maintained at 37°C and 5% CO_2_.

The viral strain used in this study was the lineage B.1 SARS-CoV-2/England/IC19/2020 (IC19) isolate (EPI_ISL_475572) [33]. All work involving the use of SARS-CoV-2 was performed in a Biosafety Level 3 (BSL-3) laboratory at St Mary’s Campus of Imperial College London. For these studies, a SARS-CoV-2 inactivated virus was generated with Ultraviolet radiation (260 - 285 nm) for 2 minutes. Loss of infectivity was confirmed by TCID_50_ test assay in Vero E6 cells. The influenza viruses (6:2 Tky/05 and 6:2 Eng/09) used in this work were rescued by reverse genetics as previously described [34]. The 6 internal genes of 6:2 Tky/05 and 6:2 Eng/09 were from avian highly pathogenic H5N1 influenza A/turkey/Turkey/1//2005 virus (Tky/05), and low pathogenic seasonal 6:2 Eng195 H1N1pdm09 (Eng/09) respectively, combined with haemagglutinin (HA) and neuraminidase (NA) genes from A/Puerto Rico/8/34 (PR8). Briefly, plasmids, encoding internal virus segments from indicated viruses and PR8 HA and NA, were co-transfected into 293-T cells alongside pCAGGs vectors expressing the polymerase and NP proteins were then co-cultured with MDCK cells. Virus stocks were grown on MDCK cells using serum free DMEM supplemented with 1 μg/mL of TPCK trypsin. Viruses were stored in −80°C and titrated on MDCK cells by plaque assay.

### Treatments

The viruses were diluted in serum-free DMEM (supplemented with 1% NEAA and P/S) to a multiplicity of infection of 0.01 or 1. The inoculum was added to macrophages, Calu-3, Caco-2 or Vero E3 cells and incubated at 37 °C for 1h. The inoculum was then removed and cells maintained as described above. 6, 24 or 72h post infection (hpi), the culture supernatants were collected and quantified by TCID_50_ assay on Vero E6 cells by the Spearman–Karber method [35] or qPCR for the SARS-CoV-2 Envelope gene (*E*). For the IFN-α experiment, macrophages were pre-treated with 1000 U/ml IFN-α (Invivogen, UK) for 24h before SARS-CoV-2 infection. For treatment with lipopolysaccharide (LPS), cells were treated with 10 μg/ml LPS (Invivogen) for 6h.

### Real-Time Quantitative Reverse Transcription PCR (qPCR)

For SARS-CoV-2 Envelope (*E*) and influenza Matrix (*M*) gene qPCR, RNA was extracted from virus supernatants using the QIAmp Viral RNA kit (Qiagen, UK) as described by the manufacturer (Qiagen, UK). qPCR was performed using the AgPath RT-PCR (Life Technologies, UK) kit on a QuantStudio^™^ 7 Flex Real-Time PCR system (Applied Biosystems) with the following primers for the *E* gene: forward: 5’-ACAGGTACGTTAATAGTTAATAGCGT-3’, reverse:5’-ATATTGCAGCAGTACGCACACA-3’,probe:FAM-ACACTAGCCATCCTTACTGCGCTTCG-BBQ. For *M* vRNA and mRNA analysis, primers and procedures were described previously [34]. A standard curve was also generated using viral RNA dilutions of known copy number to allow absolute quantification of *E* and *M* gene copies from Ct values.

Cell RNA isolation and RT–qPCR was performed using procedures described previously [36] using the 7900HT Fast Real-Time PCR System (Applied Biosystems). Primers for GAPDH, TNF-α, IP-10, IL-6, IL-8, HLA-DR, IFN-α, IFN-β, and ACE2 have been described elsewhere [37–39]. The output Ct values and dissociation curves were analysed using SDS v2.3 and RQ Manager v1.2 (Applied Biosystems). Gene expression data were normalized against the housekeeping gene GAPDH and compared with the mock controls using the comparative C_T_ method (also referred to as the 2^−ΔΔCT^method [40]. All samples were loaded in triplicate.

### Chemokine and cytokine detection

To assess the expression of cytokines in SARS-CoV-2-infected, uninfected and LPS-stimulated macrophages, we used a Proteome Profiler^™^ Human XL Cytokine Array Kit (R&D Systems, Minneapolis, MN, USA) which contained 102 different capture antibodies that were spotted on a nitrocellulose membrane, according to the manufacturer’s protocol. Immunospots were imaged with the Azure c600 Gel Imaging System (Azure Biosystems, USA) and data (Figure S1) were analysed as spot intensities using Image J (Laboratory for Optical and Computational Instrumentation, USA). For each protein, the average signal of duplicate spots was calculated, corrected for background signals, and normalized to the average signal of the membrane reference spots (relative pixel intensity).

### Statistical analyses

To determine the significance of differences between experimental groups, one-way ANOVA analysis followed by Tukey’s multiple comparisons test were carried out. *P*-values were set at 0.05 (*P* ≤ 0.05) unless indicated otherwise. Error bars represent standard deviation (SD). All data analyses and preparation of graphs were carried out with GraphPad Prism version 8.01 (GraphPad Software, San Diego, CA).

## Results and discussions

### Macrophages are refractory to SARS-CoV-2 infection

Severe infections of SARS-CoV-2 are associated with a cytokine storm characterised by high levels of IL-6 and TNF-α in patients [41]. The original source of this hyperinflammation has not yet been elucidated. Macrophages play a critical role in immune defence against virus infections and are critical for activation and resolution of systemic inflammation [42–44]. The degree to which macrophages contribute to SARS-CoV-2 propagation within the host and host immune responses to the infection is not yet clear.

We initially investigated the ability of human macrophages to support SARS-CoV-2 replication. We infected monolayers of primary human monocyte-derived macrophages (MDM) with a SARS-CoV-2 B.1 lineage strain and two recombinant influenza isolates (6:2 Tky/05 and 6:2 Eng/09) at a multiplicity of infection (MOI) of 0.01 or 1. The engineered viruses have an identical ability to bind and enter cells because they encode the same HA/NA pairing (PR8), but differ in their interaction with factors inside the infected cells depending on the human (H1N1 6:2 Eng/09) or avian (H5N1 6:2 Tky/05) virus origin of the segments encoding the internal genes and induce distinct innate immune responses [34].

We assessed productive replication of SARS-CoV-2 by titrating infectious virus in the cell supernatant at 24 and 72h post-infection (hpi) with a standard TCID_50_ assay on Vero E6 cells, and compared virus yields with those from Calu-3, Vero E6 and Caco-2. We also extracted RNA from the supernatants and the infected cells at 6 hpi and conducted qRT-PCR for the viral transcripts (*E* for SARS-CoV-2 and *M* for influenza viruses). We observed no cytopathic effect, no infectious yield and no *E* gene RT-PCR signal in the cell lysate or the cell-free supernatant of macrophages (Fig. 1A) during infection with SARS-CoV-2. In contrast Caco-2, Vero E6 and Calu-3 cell lines produced high viral loads (4 x 10^5^-5.5 x 10^5^ TCID_50_/ml) (Fig. 1A). Together these findings indicate that macrophages don’t support SARS-CoV-2 replication in line with other reports [28–31]. In contrast, influenza *M* gene was detected in cell lysates at 6 hpi, more so in 6:2 Eng/09 infected macrophages (Figure S2). However, no significant increase of the *M* gene was detected in the cell culture supernatant infected by either influenza virus at 24 or 72 hpi and no infectious virus was measured by a plaque assay. Thus, the replication of these viruses in MDM is abortive, consistent to a report that only a small subset of influenza strains can productively replicate in primary human macrophages [45].

**Fig 1.**
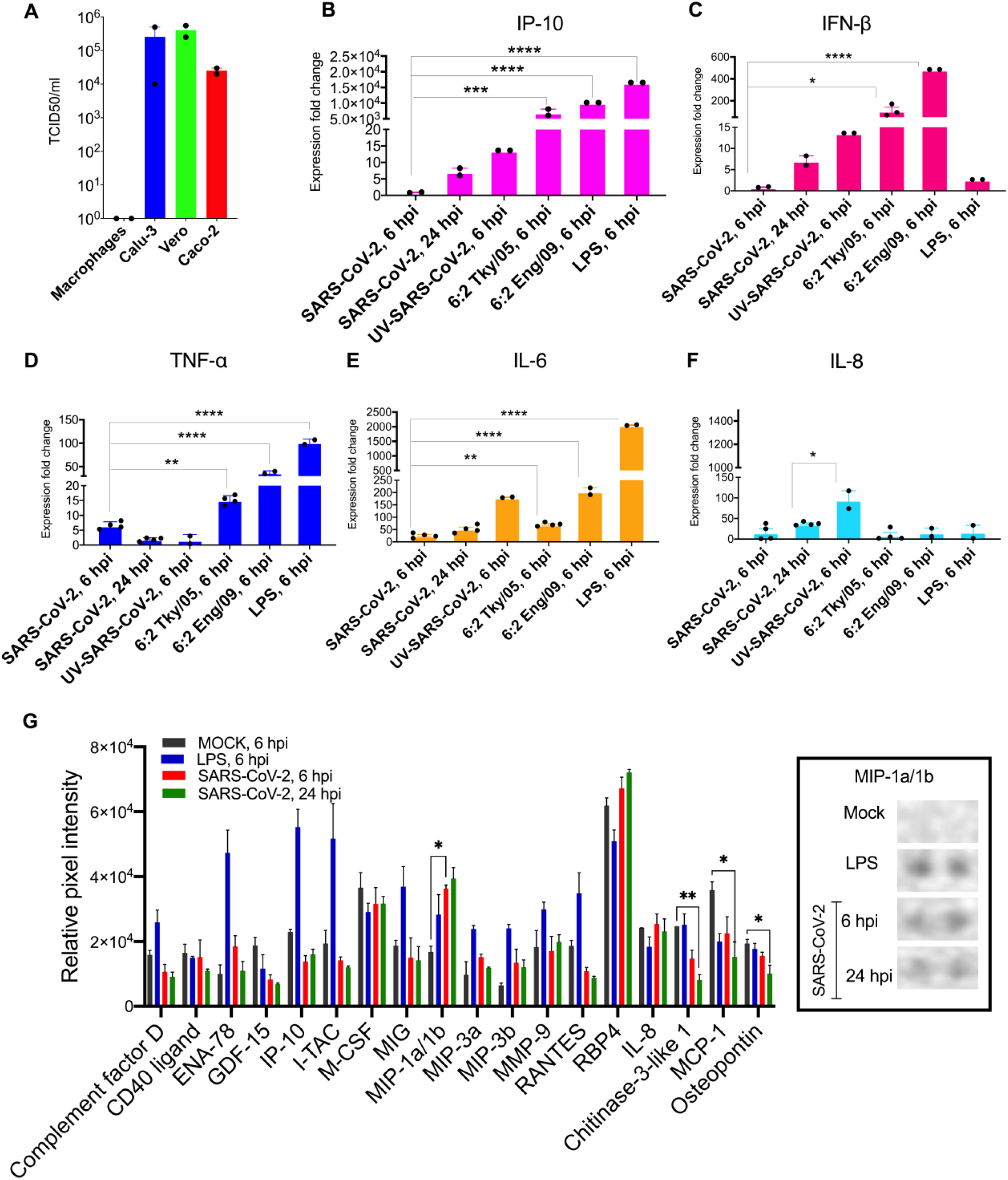
Primary macrophages are refractory to SARS-CoV-2 infection and demonstrate moderate induction of proinflammatory cytokines upon infection. (**A**) SARS-CoV-2 titres in MDM, Calu3, Vero E6 and Caco-2 at 72 hpi (MOI: 0.01) as determined by TCID50 assay. **(B-F)** The bar graphs depict the relative gene expression fold change of (**B**) IP-10, (**C**) IFN-β, **(D**) TNF-α, (**E**) IL-6 and (**F**) IL-8 mRNAs in MDM infected (MOI: 1) with SARS-CoV-2, influenza strains: 6:2 Tky/05 or 6:2 Eng/09 and MDM stimulated with LPS (10 μg/μl) compared with uninfected/untreated control. Total RNA was isolated from cells at 6 and 24 hpi. The genes were quantified by qRT-PCR. Results are presented relative to the control uninfected/untreated cell levels (2^−ΔΔ*CT*^). (**G**) (Left) The differential expression shown as mean relative pixel density and SD of 18 cytokines associated with COVID-19 screened out by the Proteome Profiler Human XL Cytokine Array kit (spotted with 102 different cytokine antibodies). The cytokine array was performed according to the manufacturer instructions using supernatant from mock, LPS-stimulated and SARS-CoV-2 infected MDM at 6 and 24 hpi (samples derived from the same assays with Fig 1B-F). (Right) Dots corresponding to MIP-1α/-1β. Statistical analysis was performed by using a one-way analysis of variance (ANOVA) followed by Tukey’s test. ***, *P*<0.001; ** *P*<0.01 *, *P*<0.05. Data derived from at least two independent experiments; means and SD are shown.

### Macrophages exhibit modest pro-inflammatory responses during SARS-CoV-2 infection

Having established that MDM did not support productive replication of either SARS-CoV-2 or influenza, we then examined the extent to which exposure of MDM to SARS-CoV-2 and influenza viruses can lead to activation and production of pro-inflammatory cytokines early in infection (6 hpi). We assessed the mRNA induction of innate antiviral and proinflammatory cytokines TNF-α, IP-10, IFN-α, IFN-β, IL-6, IL-8 and the macrophage activation marker HLA-DR in infected or lipopolysaccharide (LPS)-activated MDM. Cell response following exposure to UV-inactivated, replication-deficient SARS-CoV-2 virus was also assessed. Results are shown as expression fold change against mock MDM (Fig 1B-F). The results demonstrate that SARS-CoV-2 exposure induced significantly lower expression of IP-10, TNF-α, IL-6 mRNA compared to infection by influenza viruses or stimulation with LPS at 6 hpi. Numerically higher but still modest levels of these cytokines were detected in SARS-CoV-2 infected MDM at 24 hpi. The UV-inactivated virus retained the ability to induce IL-6 and IL-8 production (180- and 95-fold respectively). The 6:2 Eng/09 strain induced significantly more (2.5-fold) the expression of TNF-α compared to 6:2 Tky/05. Expression of IFN-α was below detectable levels in all conditions tested.

Next, we determined semiquantitatively the cytokine protein levels in the supernatants of mock (6 hpi), SARS-CoV-2-infected (6 and 24 hpi) and LPS-stimulated (hpi) MDM with a Proteome Profiler Human XL Cytokine Array. The array contains four membranes, each spotted in duplicate with 102 different cytokine antibodies. A densitometric evaluation revealed that the proinflammatory cytokines TNF-α, IP-10, IFN-γ, IL-6 and IL-8 cytokines were minimally detected in SARS-CoV-2-exposed MDM compared to mock MDM in line with the qRT-PCR results (Fig 1G). Instead, we noticed a significant induction of the Macrophage Inflammatory Protein (MIP-) 1α and 1β. MIP-1α/-1β are monocyte cytokines with inflammatory and chemotactic properties, which interact with CCR1, CCR4, and CCR5 [46] and have been found to be elevated in blood levels of COVID-19 patients (notably those admitted to intensive care units) [47]. Interestingly, the expression of three other cytokines were significantly reduced by SARS-CoV-2 (Fig 1G): monocyte chemoattractant protein 1 (MCP1), osteopontin (OPN), and chitinase 3-like 1 (CHI3L1). MCP-1 is a chemokine that attracts monocytes and basophils, but not neutrophils or eosinophils [48]. OPN is an integrin-binding glyco-phosphoprotein involved in the modulation of leukocyte activation [49] and CHI3L1 is a critical regulator of inflammation and innate immunity and a stimulator of ACE2 [50]. The levels of these cytokines are elevated in blood levels of COVID-19 patients and are associated with increased severity of disease. Our findings suggest that secretion of these cytokines are not associated with macrophages at least early in infection.

### Priming macrophages with IFNα boosts transcription of proinflammatory cytokines but does not allow productive replication

During virus infections, the majority of immune and epithelial cells produce type I interferons (IFNs: IFN-α, -β and -ω) upon sensing a virus. The IFNs do not directly kill the virus. They orchestrate a coordinated antiviral program via the Janus kinase (JAK)–signal transducers, the activators of the transcription (STAT) signaling pathway and the expression of interferon stimulated genes (ISGs), whose protein products directly inhibit virus infection [51]. SARS-CoV-2 has been reported to antagonise type I IFN responses in primary cells [52] and severe COVID-19 patients display impaired IFN-α production [53]. We explored whether pre-treatment of MDM with IFN-α can trigger an inflammatory response. MDM were pre-treated overnight with exogenous recombinant IFN-α and infected with SARS-CoV-2 for 24h as previously. Results show significantly higher expression levels of TNF-α, IL-6, IL-8 and HLA-DR in IFN-α treated cells compared to the untreated ones (Fig 2A-D) while levels of IFN-β (Fig 2E) and IP-10 (not presented) were below the detection limit.

**Fig 2.**
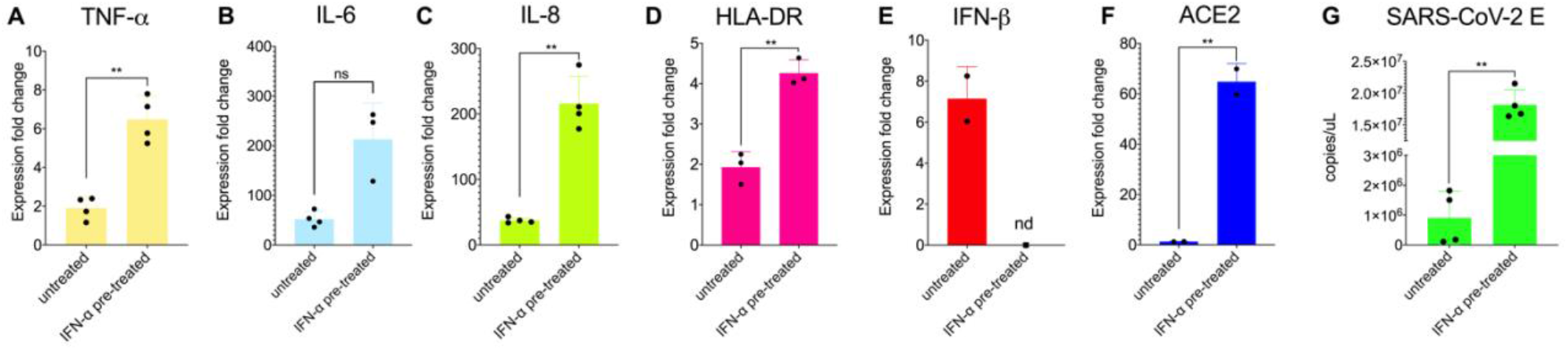
Proinflammatory response is enhanced by pretreatment with IFN-α in primary macrophages infected with SARS-CoV-2. (**A-E**) Relative fold induction of gene expression ((**A**) TNF-α, (**B**) IL-6, **(C**) IL-8, (**D**) HLA-DR and (**E**) IFN-β and (**F**) ACE2) in MDM in response to overnight pre-treatment with IFN-α (1000IU/ml) and infection with SARS-CoV-2 for 24h (MOI:1). (**G**) Quantification of subgenomic *E* RNA (qRT-PCR) in MDM infected and/or pre-treated with IFN-α. Results are presented relative to the control cell levels (2^−ΔΔ*CT*^). Statistical analysis was performed by using a paired t test. ***, *P*<0.001; ** *P*<0.01 *, *P*<0.05. Data derived from at least two independent experiments; means and SD are shown.

This finding suggests that exogenous stimuli such as secreted interferons from infected epithelial or immune cells i.e. dendritic cells, may paradoxically exacerbate the inflammatory response of macrophages in line with other reports on SARS-CoV and SARS-CoV-2 [54, 55]. We caveat that the cells we worked with here may reflect a more dominant M2 macrophage population and the observed phenotypes may differ in tissue-resident macrophages. Nevertheless, our findings are aligned with reports that macrophages are unlikely to drive the initial wave of pro-inflammatory cytokines upon SARS-CoV-2 infection [29, 31]. In addition, the substantial increase in induction of proinflammatory cytokines upon IFN-stimulation suggests that macrophages may participate in secondary or ensuing waves of inflammatory responses leading to ARDS in severe COVID-19 patients.

Interestingly, we found a significant increase of the subgenomic *E* gene in the cell lysate of IFN-treated MDM but infection was still abortive as we found no new infectious particles in the supernatant (by TCID_50_ assay and qPCR for the *E* gene; Fig 2G). Like others [56], we found that pre-treatment of MDM with IFN-α resulted in induction of ACE2 mRNA (Fig 2F). ACE2 is the receptor that SARS-CoV-2 uses to infect epithelial cells of lung alveoli and also an ISG [57]. However, a previous report that the ACE2 isoform upregulated by interferon is non-functional for SARS-CoV-2 entry [57], so our results might be explained by other changes to the cells that render them more infectable by the virus.

## Conclusions

Conflicting reports surround the role of macrophages in SARS-CoV-2 immunopathogenesis and in particular in triggering the cytokine storm that mediates the severity of ARDS in COVID-19 patients’ lungs. This study provides further evidence that macrophages are refractory to SARS-CoV-2. We report that SARS-CoV-2 failed to trigger excessive production of cytokines in macrophages at transcription and protein level, compared to two influenza viruses. We found no evidence for pro-inflammatory cytokine responses upon SARS-CoV-2 exposure. We argue that limited viral internalization or a low replication of SARS-CoV-2 in macrophages may explain the low levels of cytokines induced upon infection compared with influenza viruses. Mirroring other reports, we demonstrate that pre-treatment of macrophages with IFN-α promoted the induction of several pro-inflammatory cytokines but did not render them vulnerable to the virus despite the increased transcription of ACE2. In conclusion, macrophages are unlikely to drive the early wave of pro-inflammatory factors observed in COVID-19 patients with severe disease, but they may exacerbate immune responses later in infection.

## Abbreviations

COVID-19: Coronavirus Disease 2019
SARS-CoV: Severe acute respiratory syndrome coronavirus
TNF: tumour necrosis factor
IL: interleukin
IP: Interferon gamma-induced protein
IFN: interferon
HA: hemagglutinin
NA: neuraminidase
HPAIV: highly pathogenic avian influenza virus
ACE2: angiotensin-converting Enzyme 2
ANOVA: analysis of variance
GAPDH: Glyceraldehyde 3-phosphate Dehydrogenase

## Supplementary material

**Figure S1**: **Differential cytokine/chemokine production levels modulated by SARS-CoV-2 in MDM.** Human XL Cytokine array of secreted factors in the cell supernatant of (**A**) mock-infected, (**B**) 24 hours post LPS stimulation, (**C**) 6 hpi (MOI:1) and (**D**) 24 hpi (MOI:1) MDM.

**Figure S2: Replication of 6:2 Eng/09 and 6:2 Tky/05 influenza strains in MDM is abortive.** Quantification of *M* gene expression levels in cell lysates (red) and supernatants (blue) of MDM infected with recombinant 6:2 Eng/09 (**A**) or 6:2 Tky/05 (**B**) influenza strains (MOI:1) at 6 and 24 hpi by qRT-PCR. Bars represent Mean ± SD.

## Acknowledgements

We thank Jonathan Brown, Jie Zhou, and Thomas Peacock for their technical advice and assistance.

## Author Contributions

Z.Z. performed experiments, analysed data and wrote first draft of the manuscript; R.P. and E.S.G. performed experiments, analysed data and edited the manuscript; W.S.B. and E.S.G. conceptualized the study, supervised activities and edited the manuscript. All authors have read and agreed to the published version of the manuscript.

## Data Availability Statement

The data used and/or analysed during the current study are available, only for sections non-infringing personal information, from the corresponding author on reasonable request.

