## Supplementary material for "Primary macrophages exhibit a modest inflammatory response early in SARS-CoV-2 infection"

**Figure S1**: **Differential cytokine/chemokine production levels modulated by SARS-CoV-2 in MDM.** Human XL Cytokine array of secreted factors in the cell supernatant of (**A**) mock-infected, (**B**) 24 hours post LPS stimulation, (**C**) 6 hpi (MOI:1) and (**D**) 24 hpi (MOI:1) MDM.

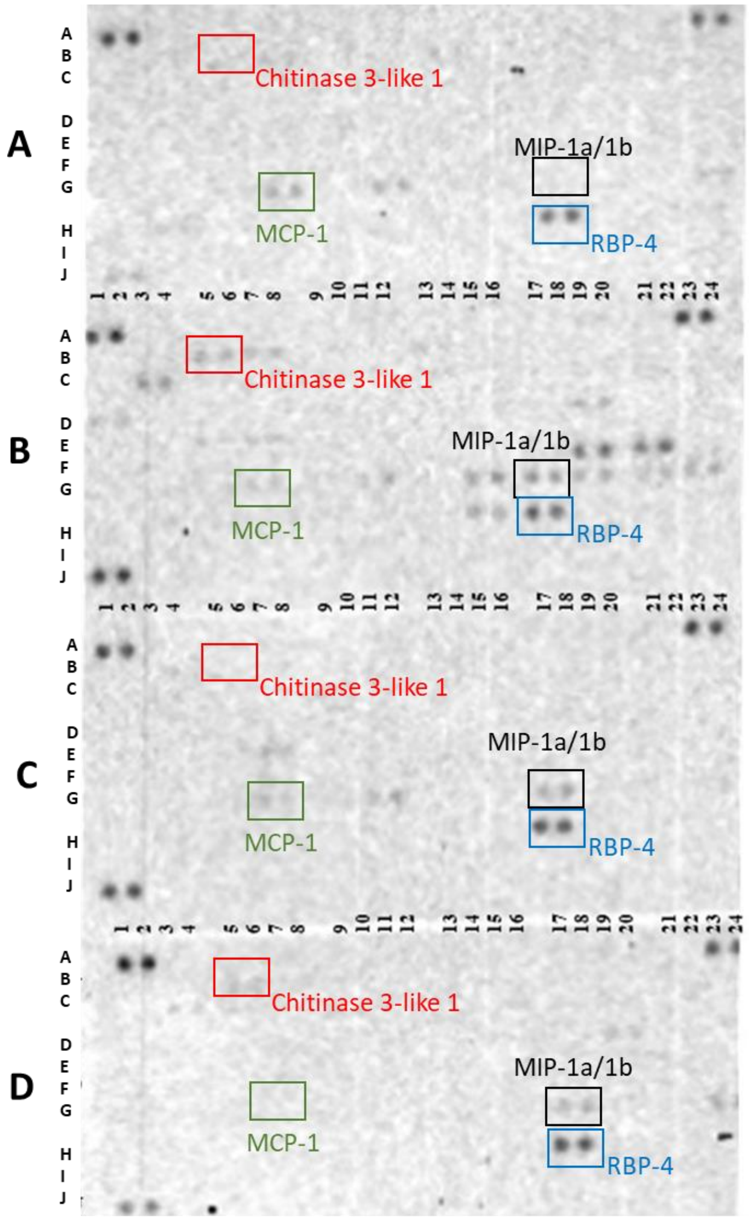

**Figure S2: Replication of 6:2 Eng/09 and 6:2 Tky/05 influenza strains in MDM is abortive.** Quantification of *M* gene expression levels in cell lysates (red) and supernatants (blue) of MDM infected with recombinant 6:2 Eng/09 (**A**) or 6:2 Tky/05 (**B**) influenza strains (MOI:1) at 6 and 24 hpi by qRT-PCR. Bars represent Mean ± SD.

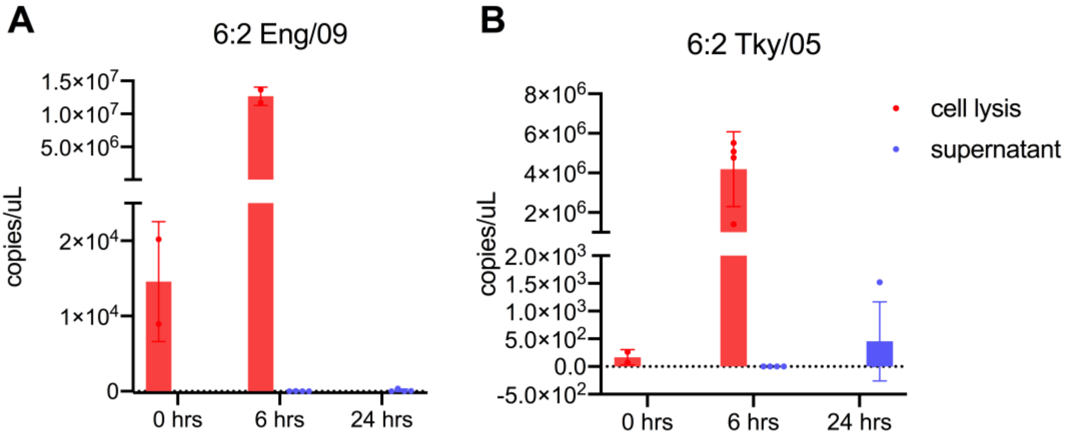
